# A microscopy-based readout to assess tumour-specific viability in neuroblastoma co-cultures and short-term cultured patient samples

**DOI:** 10.64898/2026.06.24.734168

**Authors:** Marlinde C. Schoonbeek, Sabina Valová, Sarah Swaak, Eleonora J. Looze, Merle M.J. Watzeels, Luca M. van den Brink, Mario Barrera Román, Jeroen F. van Velzen, Eoghan O’Duibhir, Karin P.S. Langenberg, Judith Wienke, Sander R. van Hooff, Marlinde L. van den Boogaard, Selma Eising, Jan J. Molenaar

## Abstract

High-risk neuroblastoma patients face poor survival despite intensive treatment. Drug testing using patient-derived models can support therapy prioritization for precision medicine and drug development. Models incorporating tumour microenvironmental components, such as co-cultures and short-term cultured patient samples containing substantial non-malignant cell fractions, could better recapitulate microenvironment-dependent drug responses. However, conventional viability assays measure the combined signal from all viable cells in a well and therefore cannot determine tumour-specific drug responses. Here, we establish a microscopy-based readout to quantify cell-type-specific viability in two complementary settings: neuroblastoma-PBMC co-cultures and freshly dissociated patient tumour samples. In the co-cultures, PBMCs were pre-labelled with a cell-tracking dye, and Calcein staining was used to independently quantify the viability of tumour cells and PBMCs in the same well. The Calcein-based viability readout correlated strongly with conventional CellTiter-Glo measurements and was compatible with automated high-throughput drug screening. The imaging workflow enabled identification of compounds with differential efficacy in co-culture versus monoculture and distinguished tumour-specific effects from PBMC toxicity. The microscopy-based viability readout was further adapted to short-term cultured patient samples. Neuroblastoma tumour cells were distinguished from the non-malignant cells using a combination of tumour-specific surface markers NCAM, L1CAM and B7H3. This enabled determination of tumour fractions and measurement of tumour-specific drug responses. Tumour fractions varied substantially between patient samples, highlighting the importance of tumour-specific viability measurements. Together, the microscopy-based viability readout for co-cultures and patient samples enables scalable assessment of tumour-specific drug responses.

## Introduction

Neuroblastoma arises from neural crest progenitor cells of the developing sympathetic nervous system and is the most common extra-cranial solid tumour in children.^1,2^ Despite intensive multimodal treatment, high-risk neuroblastoma remains associated with poor outcome, with 5-year overall survival rates of approximately 50%.^2,3^ In addition, survivors often experience severe long-term treatment-related toxicity.^4,5^ This highlights the need for more effective and better tailored treatment strategies.

Patient-derived organoids are widely used for drug testing in neuroblastoma.^6–8^ Organoid establishment usually takes several months, but once established, they are scalable for high-throughput assays which makes them well suited for drug development.^7–9^ Conventional organoid monocultures, however, lack components of the tumour microenvironment, which can influence drug response and are essential for evaluating immune-mediated therapies.^10–12^ Co-culture models incorporating immune cells increase physiological relevance while retaining scalability. However, these assays require readouts that distinguish tumour viability from immune-cell viability in the same well.^13–16^ Furthermore, drug testing on *ex vivo* short-term cultured patient material, containing unknown tumour fractions, is increasingly being explored as a complementary approach to molecular profiling for precision oncology.^8,17–20^

Most high-throughput viability assays, including CellTiter-Glo, measure the combined metabolic activity from all cells present in the well and therefore cannot distinguish tumour from non-malignant cells.^8,21,22^ Genetic labelling strategies, such as luciferase expression, can enable cell-type-specific readouts but require prior modification of the tumour cells which is time- and labour-intensive and not feasible for primary patient samples.^16^ Therefore, scalable, non-genetic, cell-type-specific viability assays are needed to assess drug responses in physiologically relevant *in vitro* models. Immunofluorescent microscopy-based approaches are increasingly compatible with high-throughput workflows and provide a solution for tumour-specific viability measurements in co-cultures and patient-derived samples.^21,23–25^ Tumour-specific markers, already established in diagnostics and therapeutic targeting, enable tumour-specific labelling in mixed cell populations. However, expression of these individual markers is heterogeneous across neuroblastoma samples, which limits the reliability of single-marker approaches.^26–29^

By combining viability dyes with either pre-labelling of PBMCs or a cocktail of tumour-specific surface markers, we establish a microscopy-based viability readout that enables separate quantification of tumour and non-tumour viability in neuroblastoma-PBMC co-cultures and patient samples.

## Methods

Detailed experimental procedures, including model establishment, culture conditions, CellTracker Orange (CTO) retention after T-cell activation, drug screening methods, imaging settings and data processing pipelines, are provided in the Supplementary Methods.

### Shared methodology

In short, two organoid lines from distinct tumour sites of the same patient^6^ were used throughout the study: AMC691T, (primary tumour; mesenchymal phenotype; HLA^+^) and AMC691B (bone marrow metastasis; adrenergic phenotype; HLA^-^).

A compound library of 134 compounds was screened in 6 concentrations in a 384-well format. Viability was normalized to vehicle-treated controls and the area under the curve (AUC) was determined from the dose-response curves. Quality control included assessment of growth rates, positive and negative controls, replicate consistency, and visual inspection of curve fits of individual curves and the overall screen.

### Neuroblastoma-immune co-cultures

In short, peripheral blood mononuclear cells (PBMCs) were obtained from healthy donors (Mini Donor Bank; institutional review board approval at UMCU: TC18-774), isolated using Ficoll density gradient centrifugation^30^ and cryopreserved at -180 °C until use.

Organoids were seeded in 384-well plates and co-cultured with PBMCs, which were pre-labelled with CTO to enable distinction between tumour and immune cells during image analysis. Co-cultures were established at an effector-to-target ratio of 1: 5 and exposed to compounds for 120 hours.

At readout, viable cells were stained using Calcein Violet (CalceinV) and imaged using the high-content confocal microscope Opera Phenix (5x air objective; confocal mode). One fixed Z-plane was selected based on comparison with a five-plane z-stack and its maximal intensity projection. CellTiter-Glo 3D (CTG) assays were performed after imaging for multiple plates, to benchmark the microscopy-based viability readout against conventional whole-well viability measurements.^7,8,22^

Image analysis was performed in Harmony software (Revvity). PBMCs were identified as CTO^+^ regions. All viable cells were defined as CalceinV^+^ regions. Viable PBMCs were then defined as overlapping CTO^+^/CalceinV^+^ regions and viable tumour cells as CalceinV^+^ regions subtracting the viable PBMC mask. The viability (CalceinV) was quantified using the sum pixel intensity of images. Compounds showing intrinsic autofluorescence that interfered with signal detection (Suppl. Table 1) were excluded from downstream analyses in the affected channel.

### Patient-derived samples

In short, patient-derived tumour samples were obtained through institutionally approved biobank protocols of the Erasmus Medical Center Rotterdam (MEC-2016-793) and the Princess Máxima Center (PMCLAB2022.370). Freshly dissociated samples were short-term cultured prior to plating in round-bottom 384-well plates. To distinguish tumour from non-malignant cells, a neuroblastoma-specific antibody cocktail (NB-cocktail) targeting NCAM, L1CAM and B7H3 conjugated to a far-red fluorophore was combined with the viability dye CalceinAM. Neuroblastoma samples (MaxPatient1-3) were cultured for 4–7 days prior to screening, whereas one melanotic neuroectodermal tumour of infancy sample (MaxPatient4) was cultured for one month until screening was performed.

Images were acquired using optimized confocal microscopy settings (10x objective; single-field acquisition; 6 z-planes) and analysed using mask-based segmentation workflows. Viable tumour cells were defined as NB-cocktail^+^/CalceinAM^+^ and total viable cells as CalceinAM^+^. The sum of the integrated pixel intensities of the CalceinAM signal was determined in both populations. Tumour fractions were calculated as the ratio of viable tumour cells to total viable cells. For proof-of-principle drug testing, patient-derived samples were exposed to selected compounds and tumour-specific viability was quantified within tumour-marker^+^ populations.

## Results

### Optimization of a microscopy-based viability readout

To enable microscopy-based measurement of viability in neuroblastoma-immune co-cultures, we first optimized and validated technical aspects of the readout method. We utilized two organoid lines derived from distinct tumour regions of the same patient: AMC691T and AMC691B (Fig. 1A).^6^ AMC691T represents the mesenchymal phenotype and expresses MHC-I (HLA^+^), enabling interaction with immune cells, such as cytotoxic T cells and natural killer cells. In contrast, AMC691B represents the more typical adrenergic neuroblastoma phenotype and lacks MHC-I expression (HLA^-^), thereby limiting immune recognition. This allowed us to model phenotypic heterogeneity regarding tumour-immune interactions while keeping other properties of the models as similar as possible.

**Figure 1:**
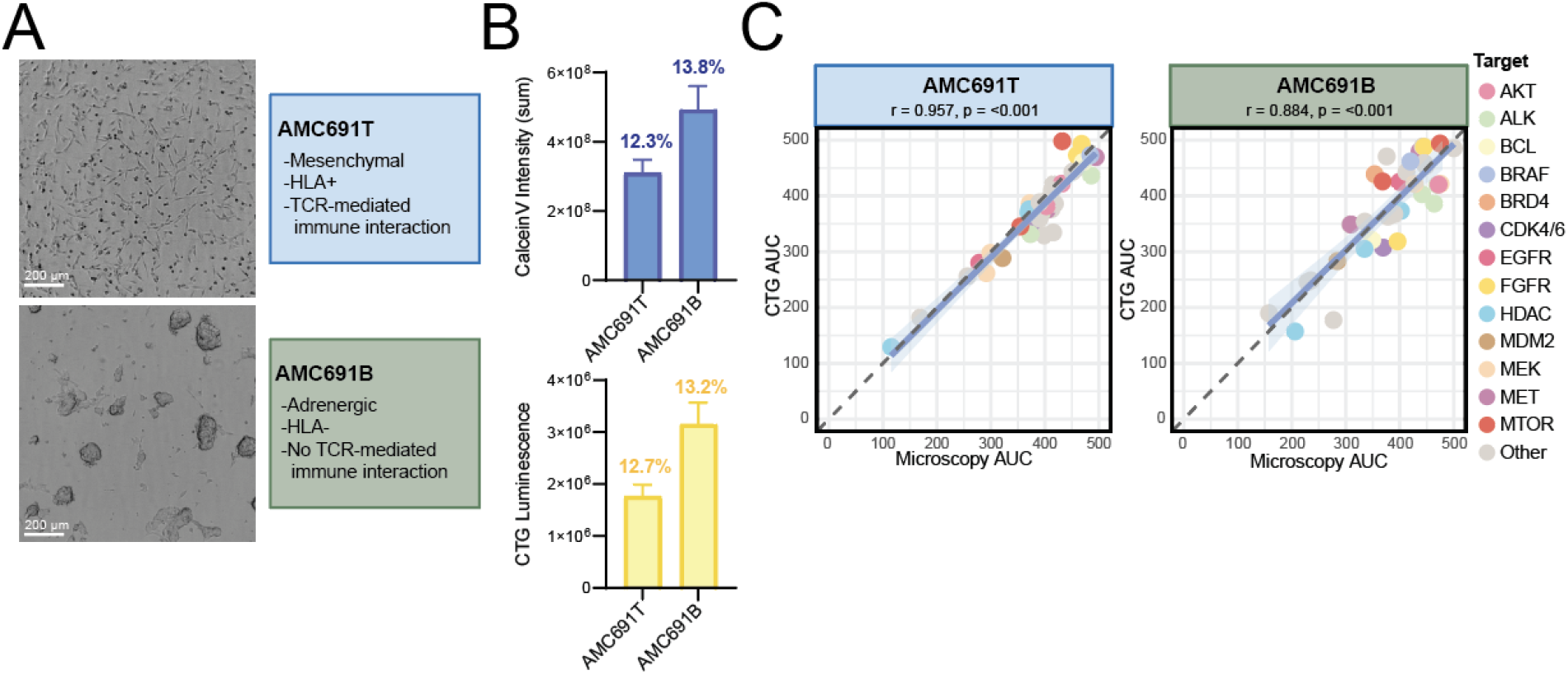
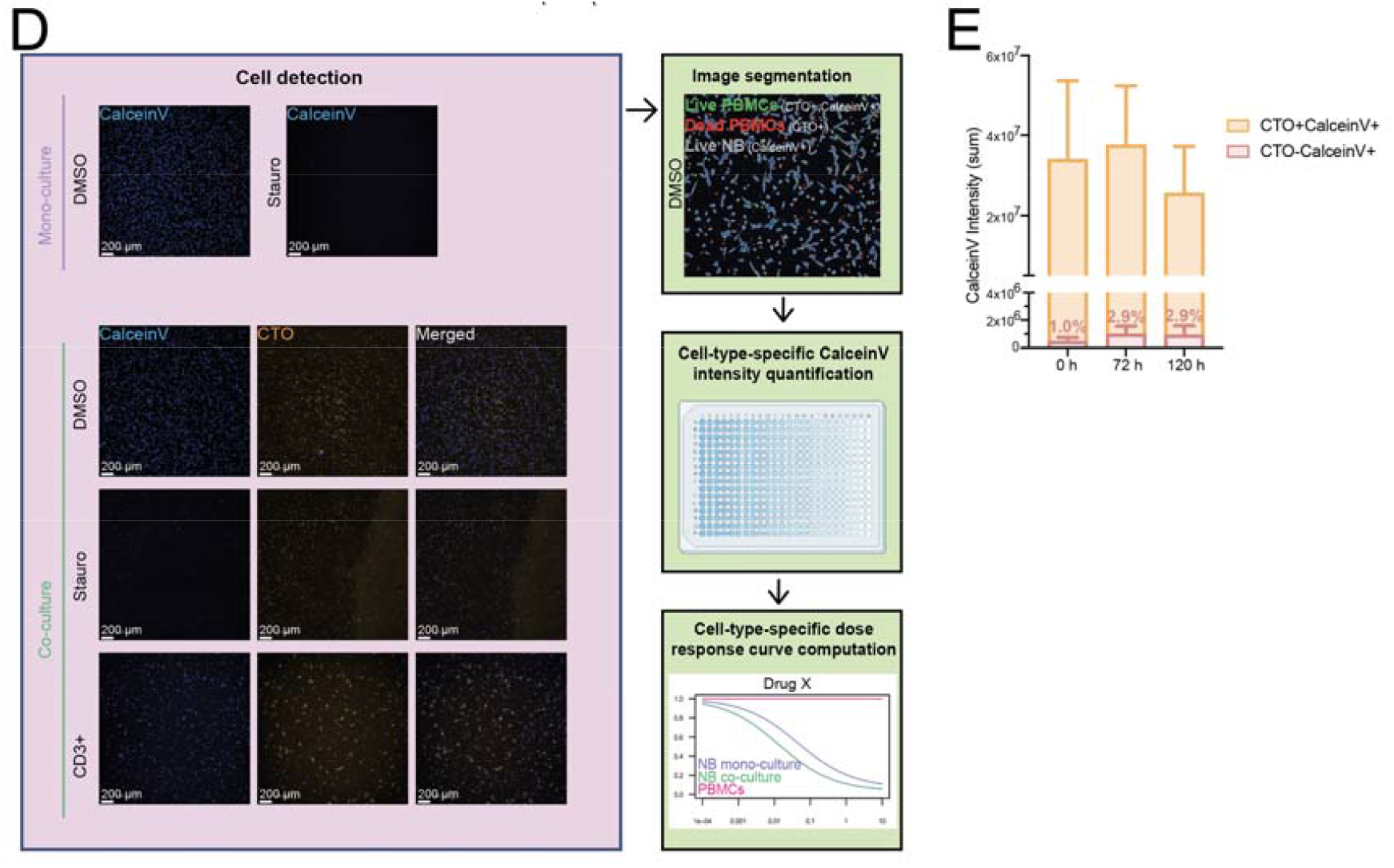
Set up and technical validation of a microscopy-based readout to reliably distinguish cell types for high-throughput drug screening in co-culture models. A) Representative microscopy images of organoid lines AMC691B and AMC691T, which are derived from different tumour locations of the same patient and differ in morphology and immune interaction. B) Bar plots representing the intensity of CalceinV (sum) and the CTG luminescence in untreated AMC691T and AMC691B organoids. The error bars represent the standard deviation. The intra-assay coefficient of variance for CalceinV (indicated above the bars) were: 13.84% (AMC691B) and 12.3% (AMC691T), and for CTG: 13.16% (AMC691B) and 12.73% (AMC691T). C) Scatter plot representing the correlation of drug response (AUC) measured by CalceinV intensity versus CTG for AMC691T and AMC691B. Each dot represents the mean AUC of two replicates normalized to the vehicle-only control of a compound (n = 45). Compounds are coloured by drug target, as indicated in the legend. Compound treatment lasted 120 hours. The Pearson correlation coefficient (r) and p-value are mentioned above the scatter plot. D) Schematic representation of the image analyses control conditions and workflow. Left part: representative microscopic images of control conditions (AMC691T): vehicle-only control (DMSO, 0.25%), positive control for cytotoxicity (staurosporine, 10 µM) and maximal immune-engagement control (CD3^+^). Images are shown for CalceinV, CTO and merged channels. Right part: analysis workflow. Images were segmented to define populations as follows: live PBMCs = CTO^+^/CalceinV^+^, dead PBMCs = CTO^+^/CalceinV^−^, and live NB = CTO^−^/CalceinV^+^. CalceinV intensity was measured within the live-NB and live-PBMC masks. Cell-type-specific dose–response curves for each compound were generated by normalizing signals to the DMSO control. E) Bar graphs representing CTO retention over time in CD3-stimulated PBMCs. PBMCs were seeded at 2,000 cells/well. The CTO^−^/CalceinV^+^ fraction indicates CalceinV signal misclassified as non-PBMCs. This fraction (indicated above the bars) was 1.04% at *t* = 0 hours, 2.94% at *t* = 72 hours and 2.89% at *t* = 120 hours. Error bars represent standard deviation. AUC = area under the dose-response curve; CTG = CellTiter-Glo 3D; CTO = CellTracker Orange; CalceinV = Calcein Violet; DMSO = Dimethyl sulfoxide; NB = neuroblastoma; PBMC = Peripheral Blood Mononuclear Cells; Stauro = Staurosporine.

To validate CalceinV as a reliable measure of cell viability, we benchmarked this imaging-based readout against the luminescence-based assay CTG, the current standard for drug screens in monocultures of organoids.^8^ Calcein Violet (CalceinV) reports on esterase activity in viable cells, whereas CTG quantifies cellular ATP content (Suppl. Fig. 1A). Microscopy-based CalceinV staining clearly distinguished viable (vehicle treated) from non-viable (staurosporine treated) cells (Suppl. Fig. 1B). The coefficients of variation in untreated wells were comparable between CalceinV and CTG signals, indicating similar assay reproducibility for both readouts (Fig. 1B; Suppl. Fig. 1C).

To compare drug sensitivities in tumour monocultures, both organoid lines were screened using a comprehensive drug library (n = 134 compounds). Comparing area under the dose response curve values (AUC), CalceinV-derived drug sensitivity correlated strongly with that of CTG (r = 0.88-0.96; Fig. 1C). Despite their phenotypic differences, overall drug sensitivity was largely conserved between AMC691T and AMC691B, with differential responses limited to a small subset of compounds (Suppl. Fig. 1D). Eight compounds, including prexasertib, showed higher efficacy (ΔAUC ≥ 100) in AMC691B (Suppl. Fig. 1E). In contrast, five compounds, including cobimetinib, were more effective in AMC691T (Suppl. Fig. 1E). Together, these results demonstrate that the microscopy-based readout provides a quantitative and robust measure of cell viability and drug response.

### Distinguishing cell type-specific viability in tumour-immune co-cultures

To assess cell-type-specific viability in co-culture conditions, peripheral blood mononuclear cells (PBMCs) from healthy donors were pre-stained with the proliferation dye CellTracker Orange (CTO) and distinguished from tumour cells using an image-based masking strategy (Fig. 1D). Downstream analysis pipelines for co-cultures segmented viable organoids as CalceinV^+^/CTO^-^ regions and viable PBMCs as CalceinV^+^/CTO^+^ regions. Cell-type specific CalceinV sum intensities were extracted from both regions. We used this approach to distinguish between viable and non-viable fractions for both cell types present in the co-culture well.

We first assessed the baseline killing of the AMC691T and AMC691B by PBMCs. Immune mediated differences were expected, since, in contrast to AMC691T, AMC691B is HLA^-^, limiting T-cell recognition. Indeed, AMC691T showed reduced viability in a PBMC co-culture compared to monoculture (Suppl. Fig. 2A), consistent with PBMC-mediated killing. To achieve ∼20% baseline killing, we tested multiple effector-to-target ratios and found 1:5 consistently optimal across three donors (Suppl. Fig. 2B). In contrast, the growth of AMC691B was comparable with and without PBMCs (Suppl. Fig. 2C).

We assessed the stability of CTO labelling over the 120-hour assay duration, including conditions of immune activation that may induce proliferation and subsequent dye dilution. PBMC monocultures were stimulated via CD3 signalling to induce TCR-mediated activation. Under these conditions, CTO loss was minimal: below 3% of viable cells misclassified as CTO^-^(Fig. 1E). PBMCs were selected for pre-labelling rather than organoids because organoids are cultured for an additional 48 hours and generally display higher proliferative capacity, potentially resulting in greater dye dilution. These results demonstrate that CTO pre-staining enables stable and reliable identification of immune cells throughout the screening assay.

Finally, to enable high-throughput screening, we automated the readout and optimized imaging speed by selecting a single Z plane, as described in the methods section. This reduced imaging time approximately two-fold, resulting in ∼8 minutes acquisition time per 384-well plate, without compromising signal intensity (Suppl. Fig. 2D-E). Collectively, these validation experiments established a robust, scalable cell-type specific microscopy-based readout suitable for high-throughput drug screening in neuroblastoma-immune co-cultures.

### Identifying compound sensitivities in co-cultures using the microscopy-based readout

Having established and technically validated this microscopy-based readout for cell-type-specific drug responses, we next applied it to high-throughput drug testing in organoid-PBMC co-cultures (Fig. 2A). In short, organoids were seeded in 384-well plates. Two days later (*t* = 0 h), pre-labelled PBMCs were added to the organoids and compounds were added for 120 hours. At readout, the viability dye CalceinV was added and automated image acquisition was performed. The drug library was screened across both conditions in the two organoid lines, AMC691T and AMC691B.

**Figure 2:**
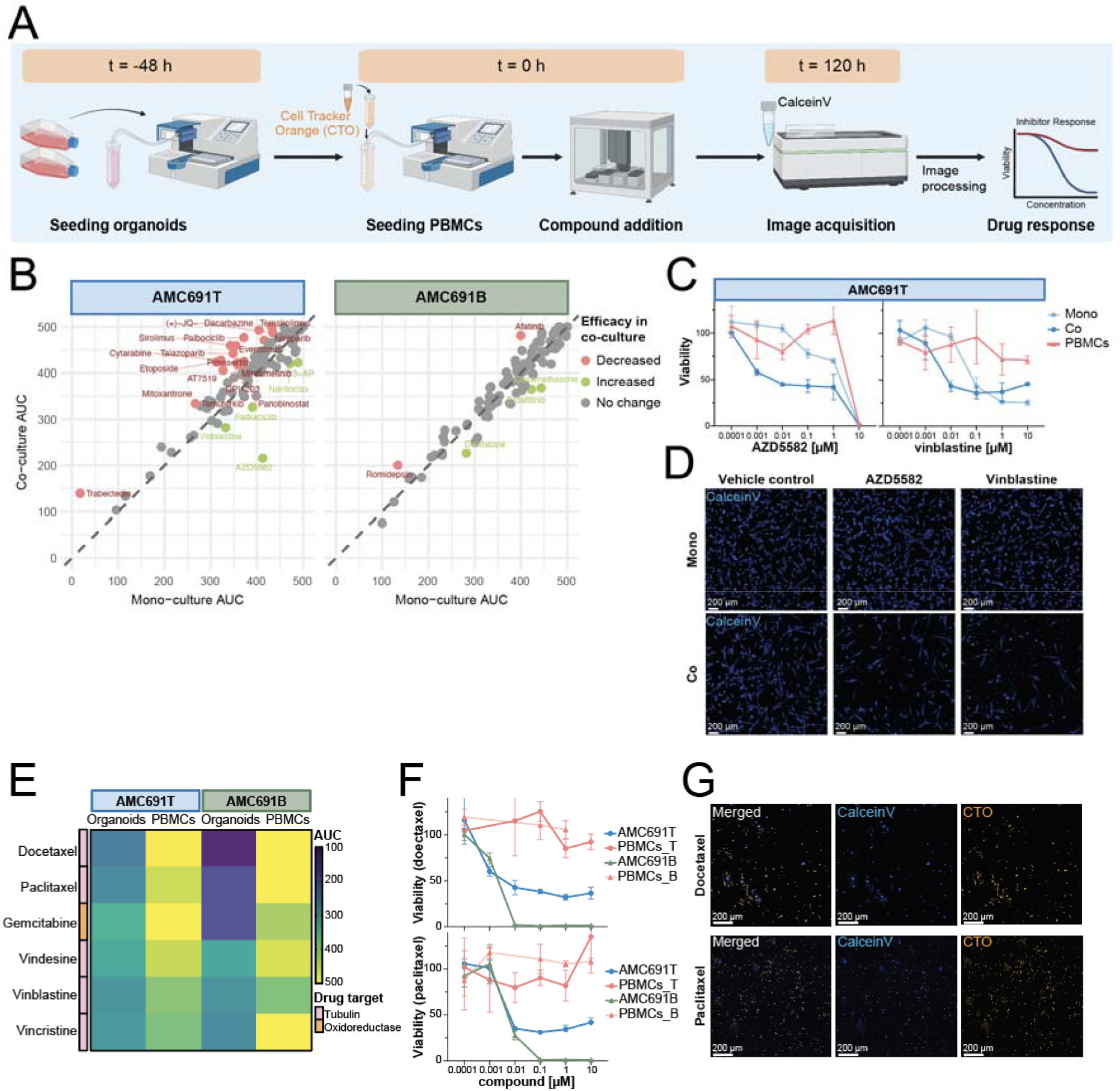
Identification of compounds with altered efficacy in mono-versus co-culture using a microscopy-based readout. A) Protocol workflow of plating, compound treatment and image acquisition. In brief, organoids are seeded (*t* = -48 hours), CTO pre-stained PBMCs were seeded and then compounds were added (*t* = 0 hours). At the day of readout (*t* = 120 hours), viability is assessed using CalceinV and images are acquired. B) Scatterplots comparing drug efficacy (AUC) of the tumour cells in mono-versus co-culture for AMC691T and AMC691B. AUC values were calculated from two technical replicates per compound and normalized to the vehicle control (0.25% DMSO) of that condition (i.e. viabilities of treated wells from the co-culture condition were normalized to the vehicle-treated co-culture condition). Each point represents one compound. Compounds with | ΔAUC| ≥ 50 were considered to show altered efficacy; red and green dots indicate decreased and increased efficacy in co-culture, respectively. C) Dose-response curves of the tumour cells of AMC691T in mono-(light blue) and co-culture (blue), and of PBMCs (red) treated with apoptosis protein antagonist AZD5582 (left) and tubulin inhibitor vinblastine (right). Symbols indicate the mean and error bars the standard deviation. AUC values are determined from this dose-response curve and represent the mean of two technical replicates (AZD5582: AUC_co = 216 versus AUC_mono = 412; Vinblastine: AUC_co = 282 versus AUC_mono = 332). D) Representative microscopy images of AMC691T in mono- and co-culture stained with CalceinV. Conditions shown: vehicle control (0.25% DMSO), apoptosis protein antagonist AZD5582 (0.001 µM); tubulin inhibitor vinblastine (0.01 µM). E) Heatmap showing drug response (AUC) for compounds with tumour-selective effects (ΔAUC ≥ 100 in organoids versus PBMCs, per line) for both organoid lines. F) Dose-response curves of organoids and PBMCs in co-culture. Colours represent AMC691T (blue), AMC691B (green) and PBMCs (red), treated with the microtubule-targeting agents docetaxel (top) and paclitaxel (bottom). Symbols indicate the mean and error bars the standard deviation. Data points exceeding the plotted y-axis range are not displayed. G) Representative microscopy images of AMC691T in co-culture treated with microtubule-targeting compounds docetaxel (0.01 µM, top) and paclitaxel (0.1 µM, bottom). AUC = Area Under the dose-response Curve; CTO = CellTracker Orange; CalceinV = Calcein Violet; DMSO = Dimethyl sulfoxide; PBMC = Peripheral Blood Mononuclear Cells

To identify compounds with altered efficacy in the presence of PBMCs, we compared drug responses between mono- and co-culture conditions (Fig. 2B). A clear distinction was observed between the two organoid models, in accordance with their immune status. Specifically, there were several compounds with altered efficacy in co-versus monoculture for AMC691T, whereas in AMC691B almost no differences were observed (Fig. 2B). In AMC691T, five compounds showed increased efficacy in the co-culture (ΔAUC ≥ 50), and 22 compounds showed reduced efficacy, in contrast to AMC691B where only 3 and 2 compounds showed increased and reduced efficacy, respectively. No overlapping compounds with altered efficacy were identified between the two organoid lines. In AMC691T, AZD5582 demonstrated greater efficacy in co-culture compared to monoculture (ΔAUC = 197; Fig. 2C-D; Suppl. Fig. 3), suggesting a PBMC-mediated effect. In line with this, AZD5582 did not exert toxic effects on PBMCs at concentrations below 10 µM (Fig. 2C). Similarly, vinblastine showed increased efficacy in co-culture (ΔAUC = 50), with PBMCs remaining viable at high concentrations (Fig. 2C-D; Suppl. Fig. 3). Together, these results demonstrate the applicability of this microscopy-based approach to identify compounds with altered efficacy in co-culture conditions, potentially informing the selection of compounds for combination with immunotherapy.

To identify compounds with tumour-selective efficacy, we directly compared tumour cell and PBMC responses from the same wells, as limiting PBMC toxicity may preserve immune-mediated anti-tumour activity *in vivo*. We identified 6 compounds that were consistently more effective in tumour cells in both organoid lines (ΔAUC ≥ 100 in organoids vs PBMCs in both lines), comprising 5 microtubule-targeting agents (docetaxel, paclitaxel, vindesine, vinblastine and vincristine) and antimetabolite gemcitabine (Fig. 2E). For example, docetaxel and paclitaxel induced tumour killing at low concentrations (0.001 µM and 0.01 µM, respectively), while PBMC viability was unaffected up to 10 µM (Fig. 2F-G). Additional tumour-selective compounds were identified in only one of the two organoid lines, with two compounds specific for AMC691T and 30 for AMC691B (Suppl. Fig. 4A). The AMC691B-specific compounds included prexasertib, decitabine, alisertib and carboplatin, which were already more effective in AMC691B monocultures and did not affect PBMC viability. Together, these results demonstrate that the microscopy-based readout enables identification of tumour-selective compounds that may be relevant for combination strategies with immunotherapies.

We next assessed compounds with comparable effects on tumour and immune cells, as the toxicity of the compounds on the immune cells may be a reason to not apply them *in vivo*. Compounds were selected based on efficacy in both cell types (AUC < 400) and limited differences in viability reduction between PBMCs and organoids (| ΔAUC| < 100). We identified six compounds, the proteasome inhibitors bortezomib and ixazomib, the HDAC inhibitors romidepsin and fimepinostat, β-catenin antagonist tegavivint and CDK12/13 inhibitor SR-4835, with similar activity in tumour cells and PBMCs across both organoid lines (Suppl. Fig. 4B-C). No compounds were consistently more toxic to PBMCs versus organoids across the full dose-response range (ΔAUC ≥ 100 in PBMCs versus organoid in both lines).

Overall, this data demonstrates the applicability of this readout for high-throughput drug screening in tumour-immune co-cultures, enabling assessment of compounds with enhanced or decreased efficacy in co-cultures, tumour-specific effects, and potential PBMC toxicity.

### Development and validation of a microscopy-based readout to determine tumour-specific viability in patient-derived samples

In short-term cultured patient material, which still contains the tumour microenvironment, it is essential to assess tumour-specific viability, similar to the co-cultures described above. To distinguish tumour from their microenvironmental cells, we developed a microscopy-based readout for use in primary patient material.

To enable drug testing in low-input patient samples, we optimized the assay for reduced cell numbers per well. Standard flat-bottom assays typically require ∼5000 cells per well, which is not feasible for testing multiple compounds on minimally invasive biopsies.^8,22^ Therefore, we adapted the assay to round-bottom 384-well plates to improve cell viability and promote spheroid growth from a limited cell density (500 cells/well). Round-bottom plates pose challenges for automated microscopy due to potential focus errors, as consecutive autofocus failures lead to termination of automated image acquisition. To address this, we optimized imaging settings using the two organoid models by systematically comparing objective magnification and numbers of field-of-view and selecting for the lowest amount of focus errors. Imaging with a 10× objective and a single central field resulted in zero focus errors, whereas imaging with multiple fields or the 5x magnification increased error rates substantially (*n* = 28 and 5, respectively; Suppl. Fig. 5A). Additionally, using only one field of view imaged 42 wells in 4 minutes versus 13 minutes for the 4 fields of view condition. Therefore, we selected the 10x objective with one central field of view, imaging the centre of the well while excluding the well edges. These optimized imaging settings enabled robust and fast image acquisition in round-bottom plates with a reduced cell input of 500 cells per well.

Patient material is more complex than neuroblastoma-PBMC co-culture systems, because tumour cells must be distinguished from multiple non-malignant cell types. As these cells cannot be selectively pre-labelled prior to plating, tumour cells were identified using a far-red-conjugated neuroblastoma-specific antibody cocktail targeting NCAM, L1CAM and B7H3 (NB-cocktail). These markers are widely used in neuroblastoma diagnostics and are being explored as therapeutic targets due to their high expression on tumour cells and limited expression in surrounding stromal and immune cells.^26–29^ By combining these markers with the viability dye CalceinAM Green (CalceinAM), it is possible to identify viable neuroblastoma cells as NB-cocktail^+^/CalceinAM^+^ and non-neuroblastoma viable cells as NB-cocktail^-^/CalceinAM^+^ (Fig. 3A).

We first evaluated the tumour markers both individually and combined on two organoid models by quantifying CalceinAM intensity within each marker-defined population. The adrenergic organoid line AMC691B showed strong positivity for all three markers, whereas its mesenchymal counterpart AMC691T was positive for B7H3 but showed minimal expression of NCAM and L1CAM (Fig 3B-C; Suppl. Fig. 5B). Combining all three markers into a single neuroblastoma-specific antibody cocktail resulted in robust detection of tumour cells in both organoids while PBMC controls showed minimal signal (Fig. 3B). We then tested the sensitivity and specificity of the NB-cocktail setup in a co-culture of AMC691B organoids with pre-labelled PBMCs and observed that the NB-cocktail distinguished tumour from immune cells (Suppl. Fig. 5C-E), with minimal overlap between the NB-cocktail and PBMC-specific eFluor450 signals. The combination of three tumour markers is particularly advantageous for patient-derived samples, where (individual) marker expression is not determined prior to the assay.

**Figure 3:**
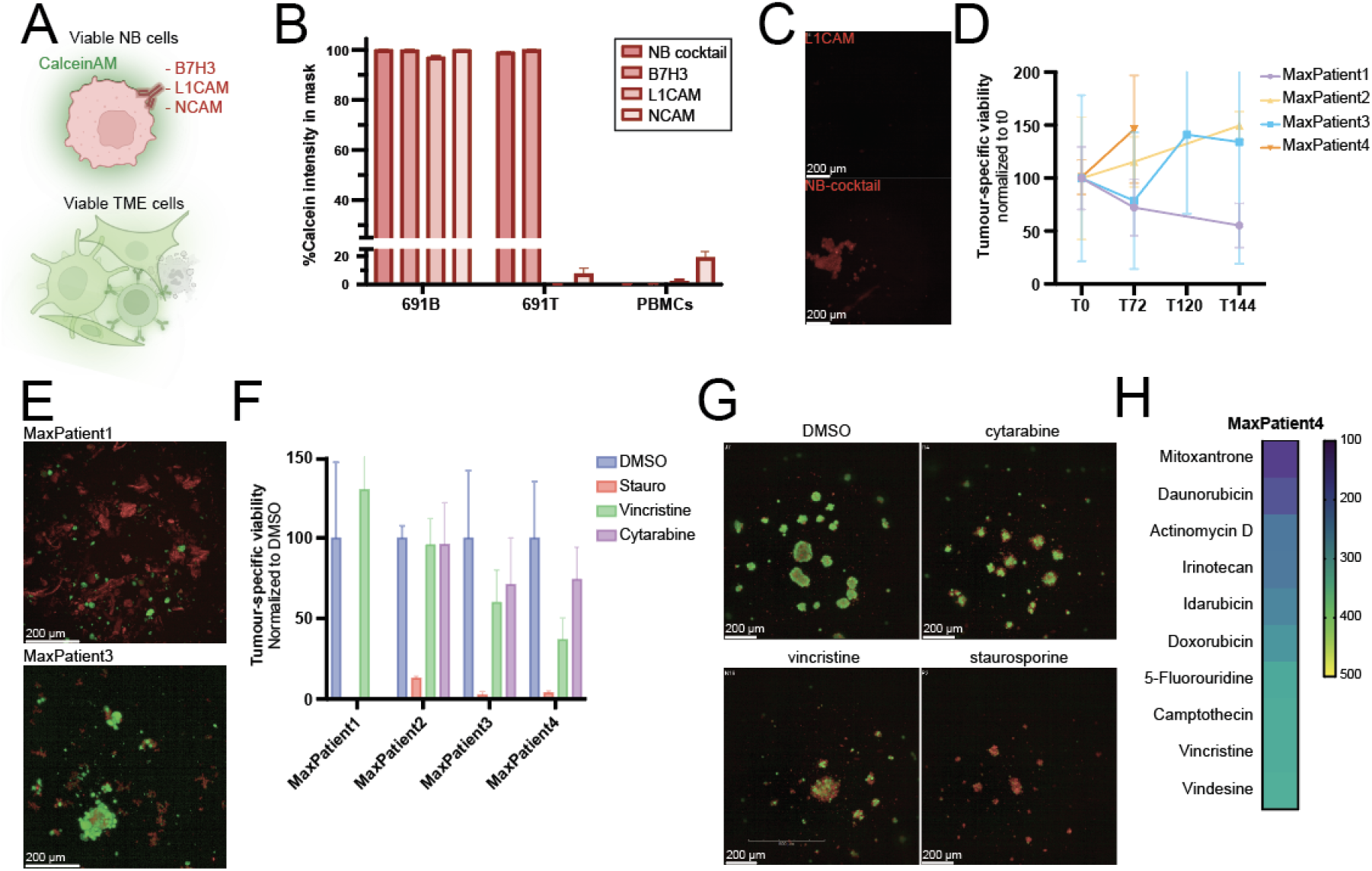
A microscopy-based readout to determine tumour fractions and tumour-specific drug responses in freshly dissociated patient samples. A) Schematic overview of the microscopy-based set up. Viable neuroblastoma (NB) cells are identified as NB-cocktail /CalceinAM and viable tumour microenvironment (TME) cells are defined as NB-cocktail /CalceinAM . The NB-cocktail consists of antibodies against the neuroblastoma-specific markers B7H3, L1CAM and NCAM. B) Bar graphs of the quantified CalceinAM sum intensity within different masks, including the NB-cocktail (1:1000 final concentration) and individual markers (B7H3, L1CAM, NCAM) for NB organoids and immune cells. Values are normalized to the total CalceinAM intensity per well. Error bars indicate standard deviation. C) Representative microscopy images of AMC691T showing L1CAM staining (top) and NB-cocktail staining (bottom; B7H3, L1CAM, NCAM). D) Graph representing tumour-specific viability, measured as CalceinAM sum intensity within the tumour-specific mask. Neuroblastoma samples were dissociated directly after tissue collection and plated within 4-7 days, the melanocytic neuroectodermal tumour of infancy sample was cultured for a month. *t* = 0 h is 24 hours after plating. E) Representative microscopy images of MaxPatient1 and MaxPatient3 representing CalceinAM (green) and NB-cocktail (red). Merged channels are shown, single channels are in Suppl. Fig. 6B. Samples were dissociated directly after tissue collection and plated within 4-7 days. *t* = 0 h, 24 hours after plating, is shown. F) Bar graphs representing tumour-specific viabilities normalized to vehicle-treated controls. Cells were treated with vehicle control (0.25% DMSO; blue), cytarabine (10 µM; purple) and MaxPatient2, MaxPatient3 and MaxPatient4 were additionally treated with staurosporine (10 µM; red) and vincristine (1 µM; green). Viability is quantified as CalceinAM sum intensity in CalceinAM^+^/tumour-marker^+^ mask and raw intensities are in Suppl. Fig. 7A. G) Representative microscopy images of neuroectodermal sample MaxPatient4 at the day of readout (*t* = 72 h), cells were treated with vehicle control (0.25% DMSO), cytarabine (10 µM), vincristine (1 µM) or staurosporine (10 µM). Merged channels (green: CalceinAM; red: B7H3) are represented, single channels are in Suppl. Fig. 7B. H) Heatmap representing tumour-specific drug response (AUC, determined by CalceinAM sum intensity in CalceinAM^+^/B7H3^+^ mask) of MaxPatient4 for the top 10 most effective compounds of the 44 tested compounds (all compounds in Suppl. Fig. 7C).

### Tumour-specific viability and drug responses in short-term cultured patient samples

To apply the established microscopy-based readout to *ex vivo* tumour material, we obtained three neuroblastoma patient samples (two biopsies and one resection) and one melanocytic neuroectodermal tumour of infancy sample (MNTI; resection; Suppl. Table 2). This rare paediatric tumour was included to demonstrate broader applicability of the workflow, since it was confirmed B7H3 positive by pathology.

We analysed the viability of the tumour and non-tumour fractions at baseline and during short-term culture. The MNTI sample (MaxPatient4) consisted almost entirely of tumour cells at *t* = 0 h (99%), whereas tumour fractions varied substantially across the neuroblastoma samples, ranging from 22% (MaxPatient3) to 66% (MaxPatient2), with MaxPatient1 at 55% (Fig. 3D-E; Suppl. Fig. 6A-B). These findings underscore the importance of tumour-specific viability measurements, as drug responses of tumour cells may be masked by large non-malignant cell fractions. Marker expression also varied between patients. MaxPatient2 showed higher expression of B7H3 and NCAM compared to L1CAM, whereas MaxPatient3 was positive for L1CAM and NCAM, but showed lower B7H3 expression (Suppl. Fig. 6C). Together, these results support the use of the NB-cocktail in patient samples with different marker expression profiles and demonstrate the applicability of the readout to determine tumour fractions and tumour-specific viability.

We next assessed whether this readout can be applied to determine tumour-specific drug responses in short-term cultured patient samples. Because of the limited amount of tumour material, we selected a small number of compounds to test at a single concentration as a proof-of-concept. Staurosporine was included as a cytotoxic compound. Vincristine was selected based on the use in standard neuroblastoma treatment regimens and, together with cytarabine, showed limited PBMC toxicity in our co-culture experiments. Due to the limited availability of viable tumour cells in MaxPatient1, only a single compound was tested. Viability was quantified within tumour-specific masks and compared with total cell viability (Suppl. Fig. 7A). The low tumour fraction at *t* = 0 h for MaxPatient3 (Fig. 3D; Suppl. Fig. 6A) was reflected by a clear difference between total and tumour-specific viability at the drug screen readout (*t* = 72 h; Suppl. Fig. 7A). In contrast, the MNTI sample MaxPatient4 showed more similar total and tumour-specific viability, consistent with the high tumour fraction. In addition, the neuroblastoma samples showed limited changes in viability during the assay (75-115% viability from 0 h to 72 h), whereas the MNTI sample did proliferate (146%).

Cytotoxicity could clearly be detected using this microscopy-based readout, as staurosporine (10 µM) induced cell death compared to the vehicle control (0.25% DMSO; Fig. 3F-G; Suppl. Fig. 7A-B). Tubulin inhibitor vincristine (1 µM) and DNA polymerase inhibitor cytarabine (10 µM) showed limited efficacy in the neuroblastoma samples, likely reflecting the limited cell proliferation (Suppl. Fig. 7A). In contrast, vincristine induced cell killing in the MNTI sample, while cytarabine showed minimal activity. Finally, the MNTI sample provided sufficient material for larger drug screen of 44 compounds across the full dose-response range (Fig. 3H; Suppl. Fig. 7C). Mitoxantrone and daunorubicin were the most effective compounds. The high tumour fraction allowed for direct comparison of CTG versus CalceinAM based drug response quantified within the tumour population, which was highly concordant (r = 0.88; Suppl. Fig. 7D). Together, this data supports the feasibility and specificity of this microscopy-based readout for the use of short-term cultured patient samples to determine the tumour fraction over time and, if sufficient material is available, to test tumour-specific drug response.

## Discussion

To advance drug testing in neuroblastoma, a difficult-to-treat paediatric cancer, we developed a microscopy-based viability readout that enables assessment of drug responses in the presence of the tumour microenvironment (TME), in both neuroblastoma-immune co-cultures and short-term cultured patient samples. This is increasingly relevant as the TME influences drug response and because immunomodulatory therapies are emerging for neuroblastoma.^10–12^ The readout strongly correlated with conventional viability measurements while providing additional value through cell type-specific viability quantification.^7,8^

The microscopy-readout for organoid-PBMC co-cultures enables direct comparison of drug effects on tumour cells and PBMCs within the same well, in contrast to approaches that assess PBMC toxicity in separate monoculture assays independently from tumour cell viability.^8,31,32^ Compounds effective against tumour cells either showed comparable PBMC toxicity or largely spared PBMC viability. The latter may preserve anti-tumour immune activity *in vivo* and thereby support combination strategies with immunotherapies.

Microtubule-targeting agents were especially effective against tumour cells and had little effect on PBMC viability. These agents disrupt microtubule dynamics, leading to mitotic arrest and apoptosis, processes to which highly proliferative cells are particularly vulnerable.^33,34^ Both organoid lines proliferated substantially more than PBMCs *in vitro*, which may partly explain their dependency on microtubule dynamics and the resulting cytotoxic selectivity. Although PBMC viability represents only a limited measure of non-tumour toxicity, this high-throughput workflow supports identification of tumour-selective compounds to be further validated in future studies.

Several compounds showed increased efficacy in co-culture compared to monoculture, including the apoptosis protein antagonist AZD5582 and microtubule targeting agent vinblastine. Previous studies have indicated immunomodulatory effects for AZD5582, including enhanced tumour infiltration by CD8+ T-cells, reduced regulatory T-cell abundance, dendritic cell activation and improved T-cell priming.^35–37^ Similarly, vinblastine has been shown to stimulate anti-tumour immune responses through activation of dendritic cells, although dendritic cells are present at low frequencies in PBMCs.^38^ These findings illustrate the value of the co-culture screen in scalable detection of compounds whose efficacy is enhanced by the presence of immune cells. Prioritized candidates can subsequently be characterized using immune profiling approaches, including evaluation of their effects on different immune cell subsets, cytokine production and activation and exhaustion markers. ^39^

The co-culture readout used allogeneic PBMCs from healthy adult donors to allow standardization, reproducibility and scalability, but these may not fully recapitulate the immune composition and dysfunction observed in neuroblastoma patients.^12,40^ Future studies could incorporate autologous patient-derived immune cells, although limited material from minimally invasive paediatric procedures remains a challenge. PBMCs represent only part of the TME and incorporating additional cell types such as stromal cells would increase physiological relevance. The workflow is also suited for studying emerging immune-mediated therapies, such as CAR-T cells, which are increasingly relevant in neuroblastoma and could be incorporated into co-culture screens.^12,41^ This co-culture workflow provides a scalable assay for tumour-immune drug screening, with potential to inform both immunotherapy development and tumour-selective compound prioritization in neuroblastoma.

We extended the microscopy-based viability readout to short-term cultured patient material to quantify tumour fraction and tumour-specific drug response in mixed cell populations. This addresses an important limitation of current *ex vivo* drug screening approaches, which rely on whole-well viability measurements and cannot distinguish tumour from non-malignant cells.^42,19^

Combining neuroblastoma-associated markers (L1CAM, NCAM and B7H3) into a cocktail enabled distinction of tumour cells from surrounding cells, compensating for intra-and interpatient variability in surface marker expression. Although these markers are widely used neuroblastoma markers, NCAM can also be expressed by immune cells (particularly NK cells) and B7H3 can be expressed by myeloid and endothelial cells in the tumour-microenvironment.^26–29,43–45^ Using a threshold-based separation limits false tumour-positive signals, as neuroblastoma cells have higher surface expression than immune cells. This microscopy-based viability readout for patient samples can be readily adapted to other tumour types by substituting tumour-specific markers.

Limited viable tumour material remains a challenge for *ex vivo* short-term patient screens.^8,19,22,42^ Therefore, we reduced the cell density per well. Neuroblastoma patient samples retained non-malignant cells during short-term culture, showing variable tumour fractions (22-66%), but tumour cells showed little to no proliferation during the assay, which likely contributed to the limited compound responses observed. Longer pre-culture durations may increase viable tumour cell numbers and proliferation rates but simultaneously change TME representation, as illustrated in the MNTI sample. After one month of culture, the MNTI sample consisted almost entirely of tumour cells with an increased proliferation rate, providing more viable tumour material for screening that enabled testing of a larger compound panel. The balance between preserving TME components and obtaining sufficient tumour material requires optimization based on the research question and available material.

Current precision oncology initiatives such as INFORM and ZERO rely on whole-well viability measurements for *ex vivo* drug testing.^19,20^ For cultures containing both tumour and non-malignant cells, tumour-specific viability assessment provides additional value by distinguishing tumour-specific drug responses from effects on surrounding non-malignant cells. The predictive value of *ex vivo* drug screening for treatment response in patients remains to be established in large paediatric cohorts. Microscopy-based tumour-specific viability readouts could contribute to this effort by enabling more precise drug response measurements, thereby ultimately supporting treatment stratification in neuroblastoma precision medicine.

## Conclusion

We developed a scalable microscopy-based readout that enables cell-type-specific drug response profiling in neuroblastoma-immune co-cultures and short-term cultured patient samples that could support future immunotherapy development and precision medicine.

## Acknowledgements

We thank all patients and their parents for providing samples that made this research possible. We appreciate the support given by the High Throughput Screening Facility and the Imaging Center of the Princess Máxima Center for Paediatric oncology. This work was partly supported by Oncode Accelerator, a Dutch national growth fund project under grant nr NGFOP220. Biorender was used to create figures 2A and 3A.

## Author contribution statement

Jan J. Molenaar: Writing - review & editing, Supervision, Conceptualization. Selma Eising: Writing - review & editing, Supervision, Conceptualization. Marlinde L. van den Boogaard: Writing - review & editing, Supervision, Conceptualization. Sander van Hooff: Writing - review & editing, Supervision, Conceptualization. Judith Wienke: Writing - review & editing, Supervision. Karin Langenberg: Recourses. Eoghan O’Duibhir: Resources, Investigation. Jeroen F. van Velzen: Resources. Barrera Román: Resources. Luca M. van den Brink: Resources. Merle M.J. Watzeels: Investigation. Eleonora J. Looze: Investigation. Sarah Swaak: Investigation, Visualisation. Sabina Valová: Writing - original draft, Visualization, Investigation, Formal analysis. Marlinde C. Schoonbeek: Writing - original draft, Visualization, Project administration, Investigation, Methodology, Formal analysis, Data curation.

