## Supplementary Figures for "A microscopy-based readout to assess tumour-specific viability in neuroblastoma co-cultures and short-term cultured patient samples"

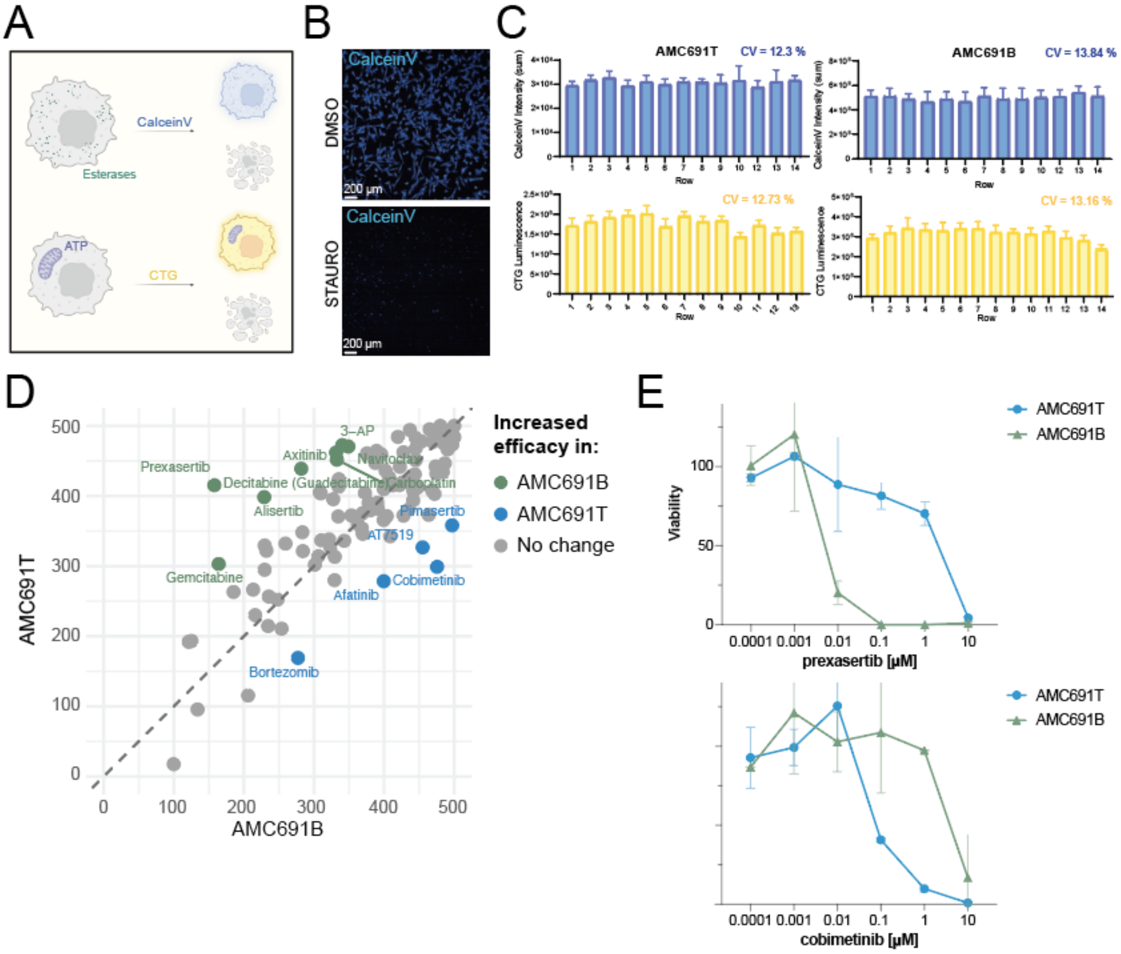


**Supplementary Figure 1: Set up and technical validation of a microscopy-based readout to reliably determine drug response in high-throughput drug screens.**

A) Schematic comparison of CalceinV and CellTiter-Glo (CTG) mechanisms. CalceinV, quantified using the sum pixel intensity of images, reports on esterase activity in viable cells, whereas the luminescence-based CTG quantifies cellular ATP content

B) Representative microscopic images of vehicle-only control (DMSO, 0.25%) and positive control for cytotoxicity (staurosporine, 10 µM).

C) Bar plots representing the intensity per row of CalceinV (sum) and the CTG luminescence in untreated AMC691T and AMC691B organoids. The error bars represent the standard deviation and intra-assay CV (%) are shown for each condition above the error bars.

D) Scatterplot comparing drug efficacy (AUC) in monoculture for AMC691T versus AMC691B. AUC values were calculated from two technical replicates per drug and normalized to the vehicle control. Each point represents one compound. Compounds with |ΔAUC| ≥ 100 were highlighted: green compounds indicate those with higher efficacy in AMC691B and blue those which were more effective in AMC691T.

E) Dose response curves of AMC691T (blue) and AMC691B (green) in monoculture, treated with CHEK1 inhibitor prexasertib (top) or MEK inhibitor cobimetinib (bottom). Symbols indicate the mean and error bars the standard deviation.

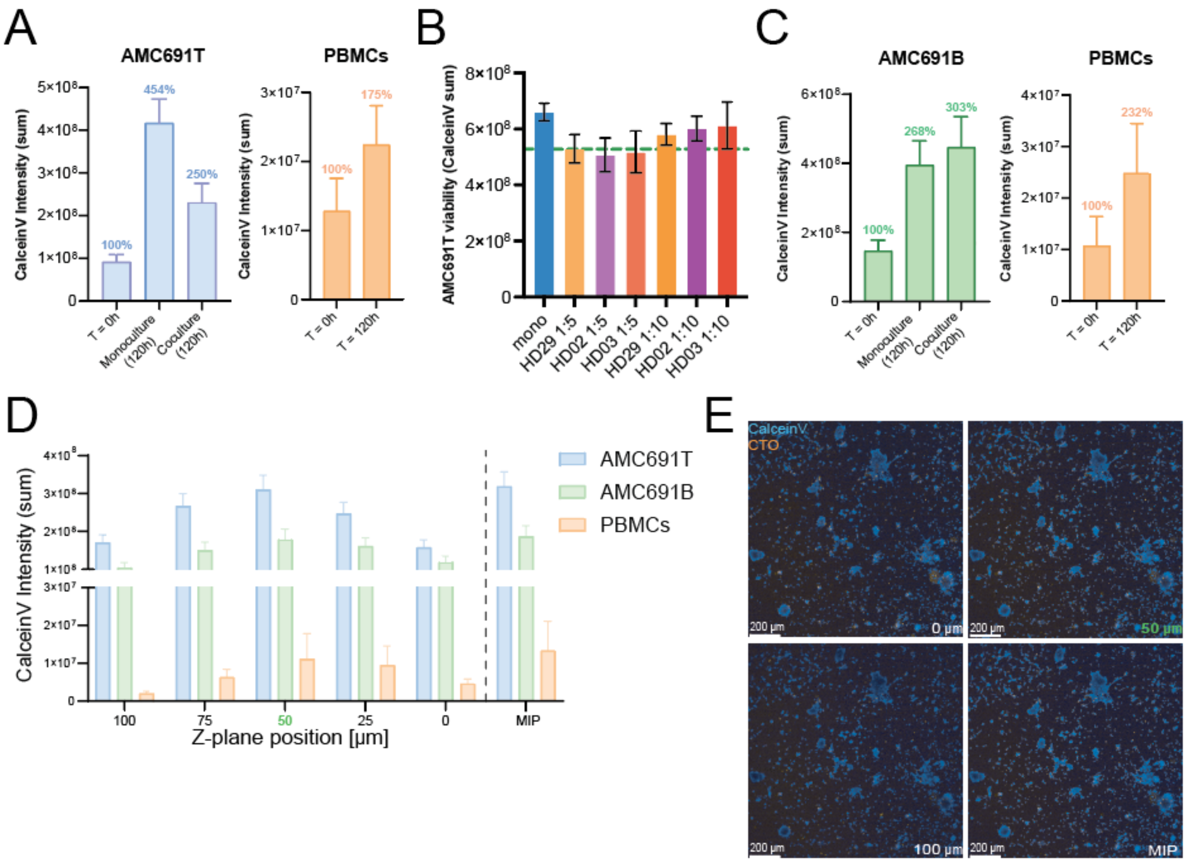


**Supplementary Figure 2: Set up and technical validation of a microscopy-based readout to reliably distinguish cell types for high-throughput drug screening in co-culture models.**

A&C) Bar plots showing CalceinV intensity of AMC691T (A) and AMC691B (C) across timepoints and culture conditions. CalceinV intensity of organoids in mono- and co-culture, and of PBMCs are shown at $t$ = 0h versus $t$ = 120h. Errors bars represent standard deviation and growth rates relative to $t$ = 0h are indicated above the bar.

B) Bar graphs representing effector to target ratios (indicated below the bars) for three different donors. Error bars represent standard deviation. The green dashed line indicates 80% cell viability, with 20% baseline killing by PBMCs.

D) Bar graph representing CalceinV intensity (sum) across individual Z-stacks versus maximal intensity projection for AMC691T/B and PBMCs. Error bars represent standard deviation. The selected Z-stack is indicated in green.

E) Representative images of AMC691B co-cultured with HD-PBMCs showing CalceinV (live cells, 450 nm) and CellTracker Orange (CTO; pre-stained PBMCs, 590 nm) across individual Z-stacks versus MIP (maximal intensity projection). The selected Z-stack is indicated in green.

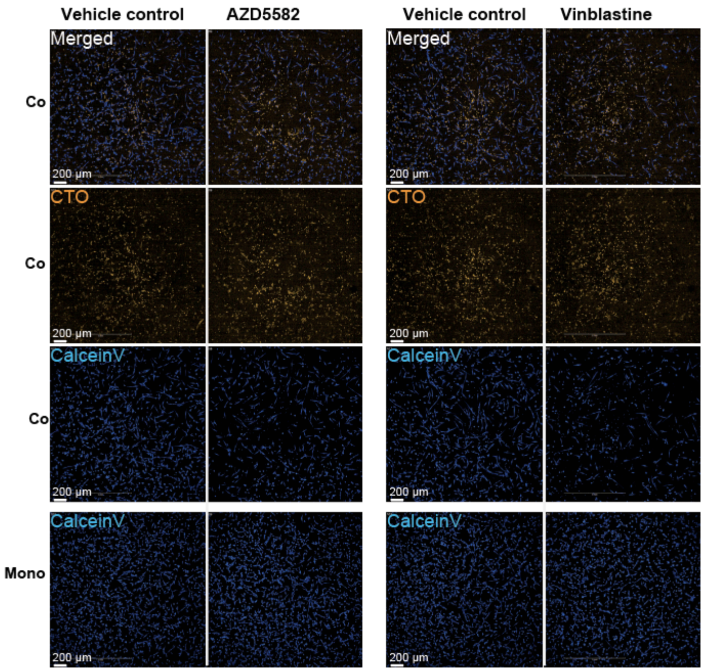


**Supplementary Figure 3: Identification of compounds with altered efficacy in mono- versus co-culture using a microscopy-based readout**.

Representative microscopy images of AMC691T in mono- and co-culture showing CalceinV, CTO and Merged channels. Conditions shown: vehicle control (0.25% DMSO), apoptosis protein antagonist AZD5582 (0.001 µM); tubulin inhibitor vinblastine (0.01 µM).

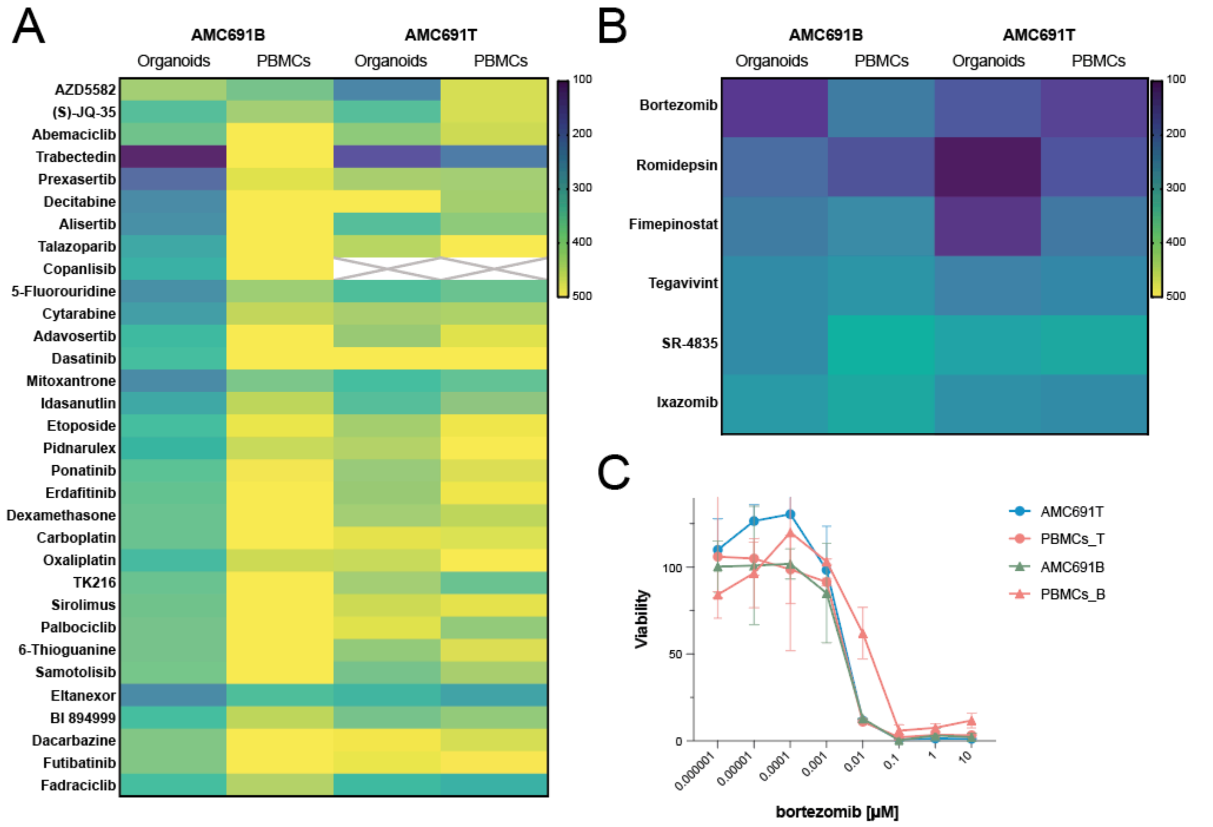


**Supplementary Figure 4: Identifying compounds with tumour- or immune- selective efficacy.**
A) Heatmap of compounds representing drug response (AUC) of compounds with tumour-selective effects. Compounds are shown for which |ΔAUC| ≥ 100 between organoids and PBMCs in one of the two organoid models. Top two compounds are based on AMC691T (AZD5582 and (S)-JQ-35), remainder on AMC691B (30 compounds).
B) Heatmap showing compounds with comparable activity in tumour cells and PBMCs for both organoid models. Compounds were selected based on efficacy towards both cell types (AUC < 400) and on a difference in AUC between organoids and PBMCs of less than 100 in both models (|ΔAUC| < 100). Values represent AUC per condition for organoid–PBMC co-cultures and PBMC-only controls.
C) Dose response curves of organoids and PBMCs in co-culture. Colours represent AMC691T (blue), AMC691B (green) and PBMCs (red), and cells were treated with the proteasome inhibitor bortezomib. Bortezomib is tested across 8 concentrations. To allow for efficacy comparison for all compounds, AUCs are determined across the 6 concentrations that are tested across all compounds (0.0001-10 μM). Symbols indicate the mean and error bars the standard deviation.

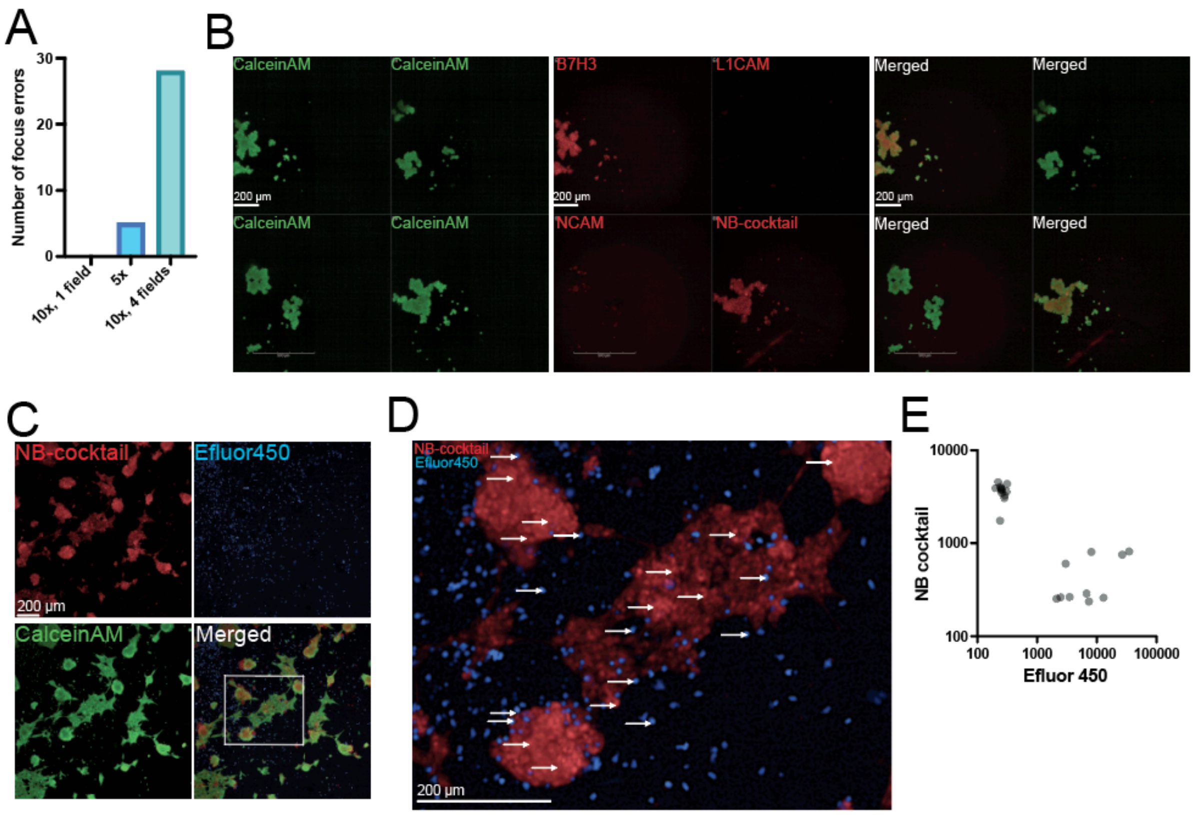


**Supplementary Figure 5: Optimization and validation of the microscopy-based readout for patient samples**

A) Bar graph representing imaging performance under different acquisition settings, measured as the number of autofocus errors per plate. The same 42 wells were imaged in different conditions. Conditions that are compared: 10x objective with 1 field of view versus the 5x objective versus the 10x objective with 4 fields of view.

B) Representative microscopy images of AMC691T stained with individual markers and the combined neuroblastoma-cocktail (NB-cocktail). Upper left: B7H3; upper right: L1CAM; lower left: NCAM; lower right: NB-cocktail (B7H3, L1CAM, NCAM). Scale bars are identical across conditions.

C) Representative microscopy image of AMC691B cells (NB-cocktail$^{+}$, red) co-cultured with pre-labelled PBMCs (eFluor450$^{+}$, blue). CalceinAM was used as a viability dye. Cells were cultured in flat-bottom plates.

D) Zoom-in of the image shown in (C), illustrating AMC691B cells (NB-cocktail$^{+}$, red) and PBMCs (eFluor450$^{+}$, blue). Arrows indicate regions for which fluorescence intensities were quantified, as shown in (E).

E) Scatter plot showing quantified fluorescence intensities for the regions indicated in (D). Each dot represents one region (n = 22).

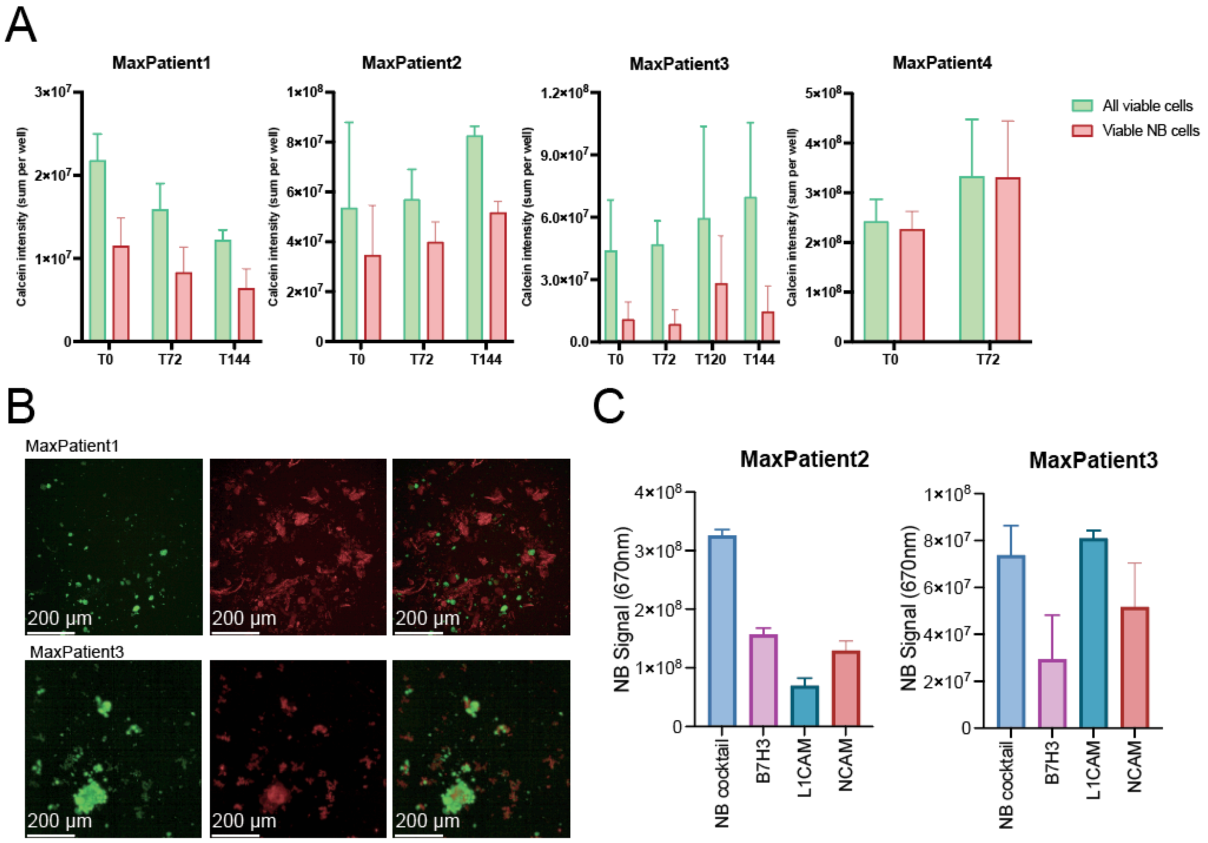


**Supplementary Figure 6: Tumour fractions of patient-derived samples in different culture conditions or over time.**

A) Bar graphs comparing the total Calcein sum intensity (green) with the Calcein sum intensity within the tumour-specific mask (red) over time for four patients. All samples were dissociated directly after tissue collection, neuroblastoma samples were plated within 4-7 days, and the melanocytic neuroectodermal tumour of infancy sample was cultured for a month. $t$ = 0h is 24 hours after plating.

B) Representative microscopy images of MaxPatient1 and MaxPatient3 representing CalceinAM, NB-cocktail. Samples were dissociated directly after tissue collection and plated within 4-7 days. $t$ = 0h, 24 hours after plating, is shown.

C) Bar graphs comparing the sum intensity of the markers in the NB+ mask of each individual marker and the combined NB-cocktail for MaxPatient2 ($t$ = 144h) and MaxPatient3 ($t$ = 72h). Each of the markers and the cocktail are added in a final concentration of 1:1000. In the NB-cocktail condition, each individual marker contributed one-third of the total antibody concentration.

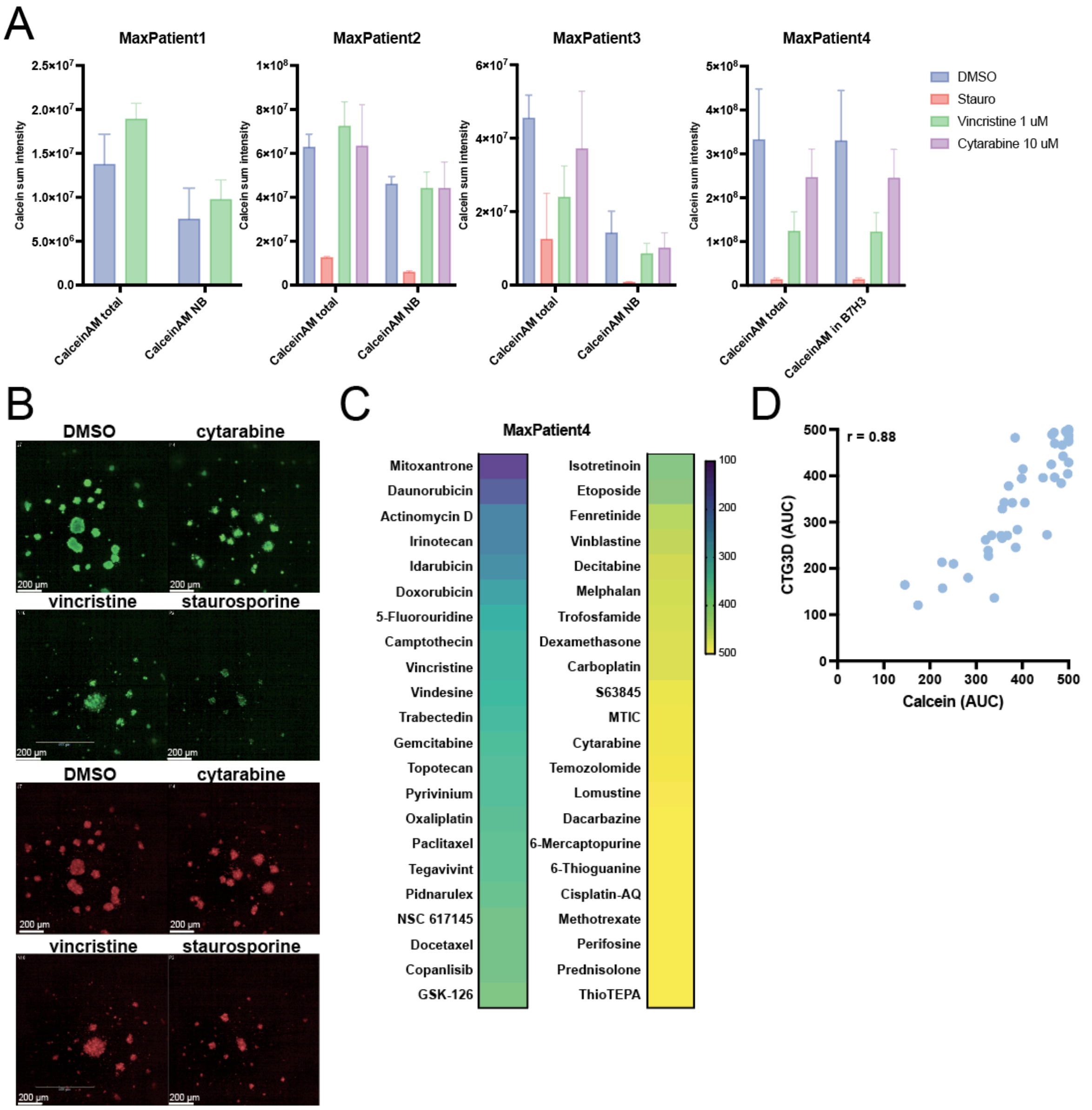


**Supplementary Figure 7: Drug testing in short-term cultured patient samples.**

A) Bar graphs representing total and tumour-specific viability (CalceinAM sum intensity in CalceinAM$^+$ mask, and CalceinAM$^+$/tumour-marker$^+$ mask) of all four patients at the day of readout (*t* = 72h). Cells were treated for 72 hours with vehicle control (0.25% DMSO; blue), staurosporine (10 µM; red), vincristine (1 µM; green) or cytarabine (10 µM; purple).

B) Representative microscopy images of neuroectodermal sample MaxPatient4 cells are treated with vehicle control (0.25% DMSO), cytarabine (10 µM), vincristine (1 µM) or staurosporine (10 µM). CalceinAM (green, top) and the NB-cocktail (red, below) are shown.

C) Heatmap representing tumour-specific drug response (AUC, determined by CalceinAM sum intensity in CalceinAM$^+$/B7H3$^+$ mask) of MaxPatient4 after 72 hours of treatment with 44 compounds, ordered by efficacy.

D) Scatterplot representing the drug response (AUC) determined by CTG or CalceinAM sum intensity in CalceinAM$^+$/B7H3$^+$ mask after 72 hours of treatment. Pearson correlation coefficient (r) is included in the top left corner.
