## Supplementary Methods for "A microscopy-based readout to assess tumour-specific viability in neuroblastoma co-cultures and short-term cultured patient samples"

**Shared methodology**

**Cell culturing**

All cultures were maintained at 37°C and 5% CO2. Patient-derived organoids were cultured in Neuroblastoma Organoid Medium (NBOM^1,2^) and passaged twice weekly. Organoids were routinely tested for Mycoplasma and authenticated by short tandem repeat profiling. AMC691B and AMC691T were established from spatially distinct tumour sites of the same patient, as previously described^3^: AMC691T, established from the primary tumour, represents a mesenchymal phenotype and expresses MHC-I on the cell surface (HLA^+^) and AMC691B, established from the bone marrow metastases, represents an adrenergic phenotype and does not present MHC-I (HLA^-^).

**Compound treatment**

Compounds from the Princess Máxima Center “Clinical Priority” library (134 compounds, clinically approved or under (pre)clinical evaluation) were added using the Echo 550 dispenser in collaboration with the High-Throughput Screening facility, as described previously.^4^ Compounds were tested at final concentrations of 0.1 nM, 1 nM, 10 nM, 100 nM, 1 µM and 10 µM (maximum of 0.25% DMSO, or MQ) and additional higher or lower concentrations were added for some compounds. DMSO-treated wells (0.25%) served as negative controls, and staurosporine (10 µM) as positive control. Anti-CD3 antibody (1:1000; soluble; ThermoFisher, #16-0037-85) was added to selected wells for maximal TCR-dependent stimulation. Cells were incubated with compounds for 120 (co-cultures) or 72 (patient samples) hours prior to viability assessment.

**Drug response analysis**

Viability was normalized to vehicle-treated controls (DMSO, defined as 100% viability). Dose-response curves were fitted using a four-parameter log-logistic model in the drc R package (version 4.0.2^5^), and the area under the dose-response curve (AUC) was determined for all compounds. Growth rates were determined by dividing the viability at day of readout by the signal at t = 0 h (>100%: growth; <100%: spontaneous cell death). Quality control included assessment of growth rates, positive and negative controls, replicate consistency, and visual inspection of curve fits of individual curves and the overall screen. AUC values were calculated across the shared concentration range (0.0001–10 µM) to enable comparison across compounds.

**Neuroblastoma-immune co-cultures**

**Isolation of Peripheral Blood Mononuclear Cells**

Peripheral blood mononuclear cells (PBMCs) were obtained from the Mini Donor Bank, as approved by the institutional review board for biobank at the University Medical Center Utrecht (TC18-774). Blood samples from healthy donors were isolated using Ficoll density gradient centrifugation as previously described.^6^ If present, residual erythrocytes were removed using red blood cell lysis (Roche, #11814389001). PBMCs were counted using Turk’s solution and a Bürker chamber and cryopreserved in FBS containing 10% DMSO (5-20 x 10^6^ cells/ mL) at -180 °C until use.

**Neuroblastoma-PBMC co-culture**

At *t* = −48 h, organoids were dissociated to single cells using Accutase, counted using an automated cell counter (TC20, Bio-Rad) and seeded into 384-well plates (Revvity, ViewPlate; #6007460) at 2000 cells per well in 35 µL NBOM using the MultiDrop Combi Reagent Dispenser (ThermoFisher).

At *t* = 0 h, PBMCs were thawed, stained with CellTracker Orange (CTO; ThermoFisher; #C7025; 1:1000 dilution) for 20 minutes at 37 °C in the dark and washed. PBMCs were added to organoids in an effector-to-target ratio of 1:5, taking the tumour growth during the 48-hour pre-culture into account, in 35 uL lymphocyte medium (RPMI supplemented with 10% FBS, 1% penicillin-streptomycin, and 1% L-glutamine). Monocultures received equal volumes of lymphocyte medium. Co-culture plates contained 16 vehicle-treated wells of only organoids in absence of PBMCs. The same healthy donor was used for drug screening of AMC691B and AMC691T to ensure comparison. The selected effector-to-target ratio yields approximately 20% tumour killing in HLA+ organoids, allowing for sufficient dynamic range for synergy with tested compounds.

**Image acquisition to assess cell viability**

At readout (*t* = 120 h), Calcein Violet 450 AM (CalceinV; ThermoFisher; #65-0854-39; 1.7 µM) was added using the Echo 550 dispenser. Plates were centrifuged at 100 g for 1 minute to ensure even dye distribution across the plate prior to incubation at 37 °C for 50 minutes, followed by 10 minutes equilibration inside the Opera Phenix microscope chamber (set to 37 °C and 5% CO₂) prior to imaging. Additionally, for both co-cultures and mono-cultures, a readout at *t* = 0 h is performed on an extra plate to determine cell growth during the screen and to assess baseline immune-mediated effects.

Images were acquired using a 5x air objective with double-peak autofocus at a fixed Z-position in confocal mode. A single Z-plane was selected, based on comparison with a five-plane Z-stack each 25 µm apart and its maximal intensity projection, which showed comparable CalceinV signal intensities for both tumour cells and PBMCs. Acquisition settings for the CalceinV channel were: excitation 405 nm 435-515 nm emission filter, 50% laser power and 60 ms exposure time. For the CTO channel: excitation 561 nm, 570-630 nm emission filter, 50% laser power, and 40 ms exposure time were applied.

For selected plates, a Cell Titer Glo 3D (CTG, Promega, #G9683) cell viability assay was performed post-imaging using the supplier’s protocol, to benchmark viability measurements as this is the conventional readout for neuroblastoma organoids in monoculture and for drug screens on short-term cultured primary (patient) material.^1,2,4^

**Image analysis to obtain tumour-specific viability from co-cultures**

Image analysis was performed in Harmony software (Revvity, version 5.3). PBMCs were identified as CTO^+^ regions (function: Find Image Region) with an absolute threshold between 500- and 5000-pixel intensity values (capturing PBMC signal while excluding any background and bleed-through), using the unstained control to set the threshold. All viable cells were defined as CalceinV⁺ regions (function: Find Image Region with Common Threshold method (≥0.34)) using staurosporine-treated positive controls for thresholding. Viable PBMCs were then defined as overlapping CTO^+^/CalceinV^+^ regions (function: Select Population, >50% overlap inclusion criterion) and viable tumour cells as CalceinV^+^ regions subtracting the viable PBMC mask (function: inverted mask, >50% overlap). The sum of the integrated pixel intensities of CalceinV was quantified within both masks to assess separate viabilities for both populations. Compounds showing intrinsic autofluorescence that interfered with signal detection (Suppl. Table 1) were excluded from downstream analyses in the affected channel.

**Anti-CD3 stimulation**

PBMCs were stained with CTO and stimulated with soluble anti-CD3 antibody (1:1000) to induce T cell receptor engagement and T cell proliferation. PBMCs were seeded into 384-well plates at 2000 cells per well (drug screen condition) and CalceinV intensity was measured in both CTO+ and CTO− populations to determine the extent of CTO dilution during proliferation.

**Primary samples**

**Sample processing**

Dutch tumour samples were obtained through an institutionally approved research study by the institutional review board of the Erasmus Medical Center Rotterdam (MEC-2016-793), and the biobank committee of the Princess Máxima Center (PMCLAB2022.370). Tumour samples were processed within 4 hours after surgery. Samples were dissociated mechanically and enzymatically if needed, using Collagenase IV for maximal 60 minutes at 37°C, and short-term cultured prior to plating. Neuroblastoma samples (MaxPatient1-3) were cultured for 4-7 days and the melanotic neuroectodermal tumour of infancy (MNTI) sample (MaxPatient4) was cultured for one month until screening was performed.

After short-term culture, patient samples were filtered (70 µm cell strainer) and manually counted (Trypan Blue and Bürker counting chamber). A neuroblastoma-specific antibody cocktail (NB-cocktail) was added to the medium (1:1000 final concentration) prior to plating. The NB-cocktail consists of anti-B7H3 (CD276 APC-M; ThermoFisher, #7-517), anti-NCAM (NCAM CF-640R; Biotium, #BNC400795-100) and anti-L1CAM (L1CAM CF-640R; Biotium, #BNC400679-100) which are all conjugated to a far red fluorophore (APC-M or CF-640R). If sufficient material was available (MaxPatient2 and 3), single stains of the markers were also plated (1:1000 final concentration). Cells were plated manually or using the Multidrop dispenser (MaxPatient4) at 500 cells / well in round-bottom 384 well plates (Corning; 3830) in 40 µl NBOM containing the NB-cocktail. CalceinAM (ThermoFisher; #C34852; 0.5 µM) was added at readout as described in *Image acquisition to assess cell viability.*

**Image acquisition**

Imaging was performed in confocal mode using a 10x objective with single-field acquisition, one peak autofocus and 6 z-planes. Autofocus settings were optimized to minimize acquisition failure, as consecutive focus errors (>5) terminate automated imaging. Plates were centrifuged to ensure consistent CalceinAM distribution prior to CalceinAM incubation at 37°C (50 min + 10 min equilibration in the microscope). Acquisition settings for the CalceinAM channel (excitation 488 nm) were: 500/550 nm emission filter, 10% laser power and 40 ms exposure time. For the NB-cocktail channel (excitation 640 nm): 650/760 nm emission filter, 100% laser power, and 200-240 ms exposure time were applied. Viable tumour cells were defined as CalceinAM^+^/NB cocktail^+^ and total viable cells as CalceinAM^+^ and the sum of integrated pixel intensities of the CalceinAM signal was determined in both populations. Tumour fraction was calculated as the ratio of viable tumour cells to total viable cells.

As we used one field of vies, in validation experiments, in some occasions on the outer columns cells had shifted towards the well edges after centrifugation. If so (checked with the DMSO-treated controls), these wells were excluded from the downstream analysis. Hence, all patient samples were plated in the middle of the 384-well plates.

As image acquisition was performed using a single central field of view, cell distribution was assessed in vehicle-treated control wells. During assay validation, in occasional wells at the outer edges of the plate, cells shifted towards the well edges following centrifugation and were therefore not fully captured. These outer columns were then excluded from downstream analysis. To minimize this effect, patient samples were plated in the central columns of the 384-well plates.
